# Working memory implements distinct maintenance mechanisms depending on task goals

**DOI:** 10.1101/162537

**Authors:** Johannes J. Fahrenfort, Jonathan Van Leeuwen, Joshua J. Foster, Edward Awh, Christian N.L. Olivers

## Abstract

Working memory is the function by which we temporarily maintain information to achieve current task goals. Models of working memory typically debate where this information is stored, rather than how it is stored. Here we ask instead what neural mechanisms are involved in storage, and how these mechanisms change as a function of task goals. Participants either had to reproduce the orientation of a memorized bar (continuous recall task), or identify the memorized bar in a search array (visual search task). The sensory input and retention interval were identical in both tasks. Next, we used decoding and forward modeling on multivariate electroencephalogram (EEG) and time-frequency decomposed EEG to investigate which neural signals carry more informational content during the retention interval. In the continuous recall task, working memory content was preferentially carried by induced oscillatory alpha-band power, while in the visual search task it was more strongly carried by the distribution of evoked (consistently elevated and non-oscillatory) EEG activity. To show the independence of these two signals, we were able to remove informational content from one signal without affecting informational content in the other. Finally, we show that the tuning characteristics of both signals change in opposite directions depending on the current task goal. We propose that these signals reflect oscillatory and elevated firing-rate mechanisms that respectively support location-based and object-based maintenance. Together, these data challenge current models of working memory that place storage in particular regions, but rather emphasize the importance of different distributed maintenance signals depending on task goals.

**Significance statement (120 words):** Without realizing, we are constantly moving things in and out of our mind’s eye, an ability also referred to as ‘working memory’. Where did I put my screwdriver? Do we still have milk in the fridge? A central question in working memory research is how the brain maintains this information temporarily. Here we show that different neural mechanisms are involved in working memory depending on what the memory is used for. For example, remembering what a bottle of milk looks like invokes a different neural mechanism from remembering how much milk it contains: the first one primarily involved in being able to find the object, and the other one involving spatial position, such as the milk level in the bottle.

## \Body

In recent years, the hypothesis that sensory areas are recruited when maintaining information in working memory (WM) has gained considerable prominence. Indeed, many studies have now shown the involvement of sensory regions (1-3), as well as spatial attention (4) during WM retention. However, while this has elucidated the role of storage, it has overshadowed the fact that an important function of WM is goal maintenance (5, 6), as WM is only useful in light of current task requirements. The importance of goal maintenance is in line with the traditionally assumed involvement of frontal cortex (7), as well as the observed coupling between memory representations and motor plans (8, 9), including the finding that maintaining information affects associated action systems (10-12). Moreover, there has been much emphasis in on the locus of storage (5, 13), underemphasizing the potentially distributed nature of working memory (14, 15) and the neural mechanisms involved in maintenance.

The current study shows with high temporal resolution how identical sensory input is transformed into dissociable distributed neural maintenance signals, depending on the requirements of the task as a whole. Twenty participants were tested in a repeated measures design in which 64-channel electroencephalograms (EEG) were obtained. Participants maintained an oriented bar across a retention interval in all conditions, but were given different tasks at test. In one task, they indicated the orientation on a continuous scale, by clicking on the point at which the bar intersected with the surrounding circle (*continuous recall task*, see Fig. 1a). In a second task, they selected the remembered orientation from a circular array of oriented bars (*visual search task*, see Fig. 1b).

**Figure 1.**
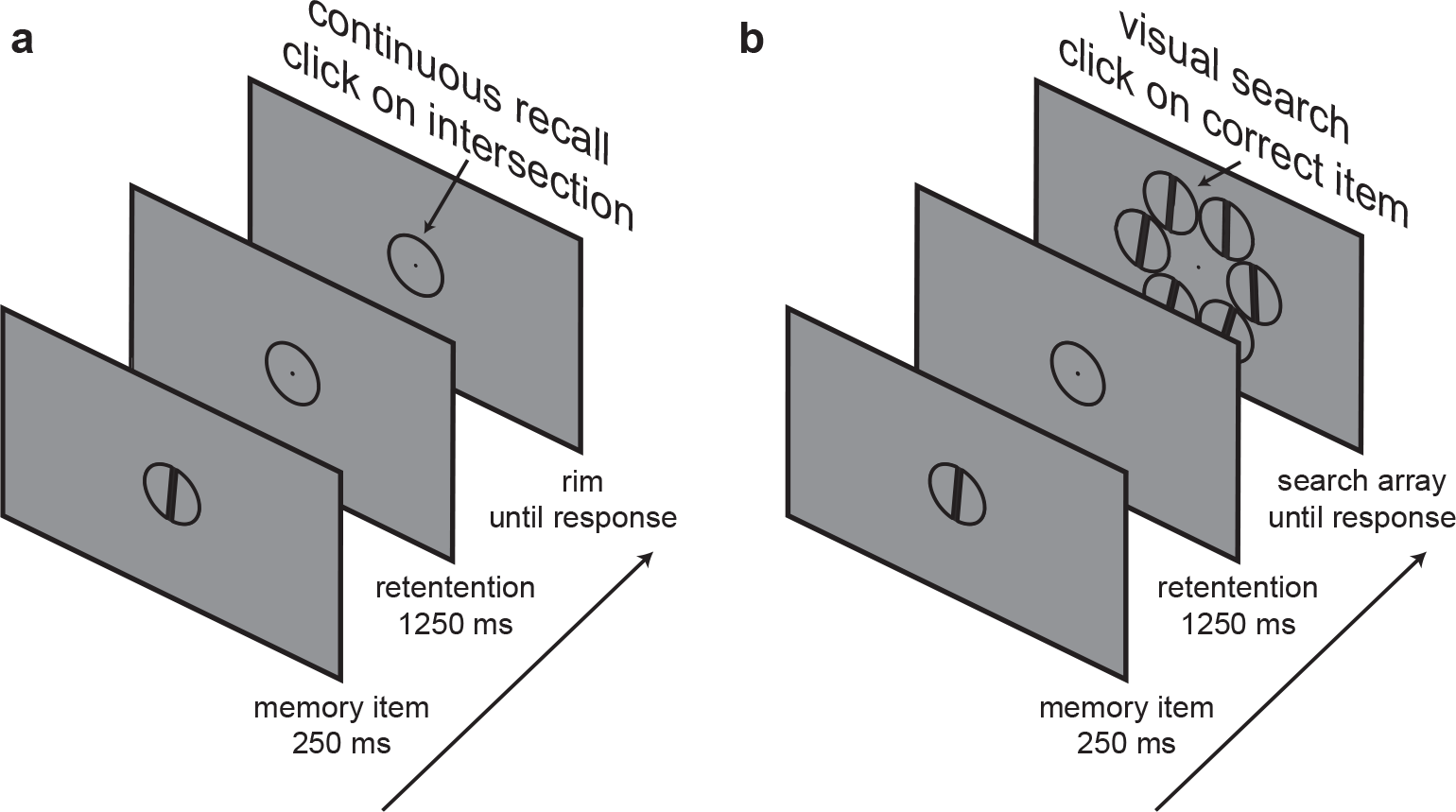
Experimental design. The initial stimulus sequence was identical, only the response screen was different between tasks. **(a)** Continuous recall task: subjects had to reproduce the item at test by clicking on the position of the circle where the bar would have intersected with the rim. **(b)** Visual search task: at test, subjects had to click on the right target item in a 6-item circularly organized search array.

The sensory input, memoranda, and temporal structure of the trial were thus identical in both tasks. Both tasks also required highly similar motor responses, by moving the mouse to a position on a circle. The crucial difference between the two tasks was that in the continuous recall task, there was a direct coupling between the response and the motor action, encouraging participants to maintain the position to which they should move the mouse to reproduce the orientation. In the search task, there was no such relationship, as the position of the target in the search array was unpredictable and unrelated to the memorandum itself. Here the task favored a mechanism allowing the maintenance of object identity. Hence both tasks were identical, including equivalent motor actions at test time, but the continuous recall task favored maintenance to be subserved by a location-based code, while the search task favored object-based maintenance. Tasks were administered in separate sessions, using a counterbalanced order across subjects.

The bar was oriented at one of 24 equidistant angles spanning 180 angular degrees to cover all spokes in a wheel. For the decoding and forward modeling analyses, these were grouped into 6 classes, each containing 4 adjacent orientations (see online Methods). To investigate the neural correlates associated with maintenance of content in the two tasks, we used a three-step analysis pipeline. First we used a backward decoding approach, applied to two potential sources of multivariate activity that could support memory in the retention interval: untransformed raw electrophysiological signals (16, 17) and time-frequency decomposed power modulations of the signal, specifically in the 10-12 Hz alpha range (18-20). Using a 20-fold train-test procedure, we first decoded class membership based on alpha-band power (Fig. 2a, see Supplementary Fig. S1a for the full time-frequency domain from 2-30 Hz) and raw EEG (Fig. 2b) using generalization across time (GAT) matrices of decoding accuracy (21).

**Figure 2.**
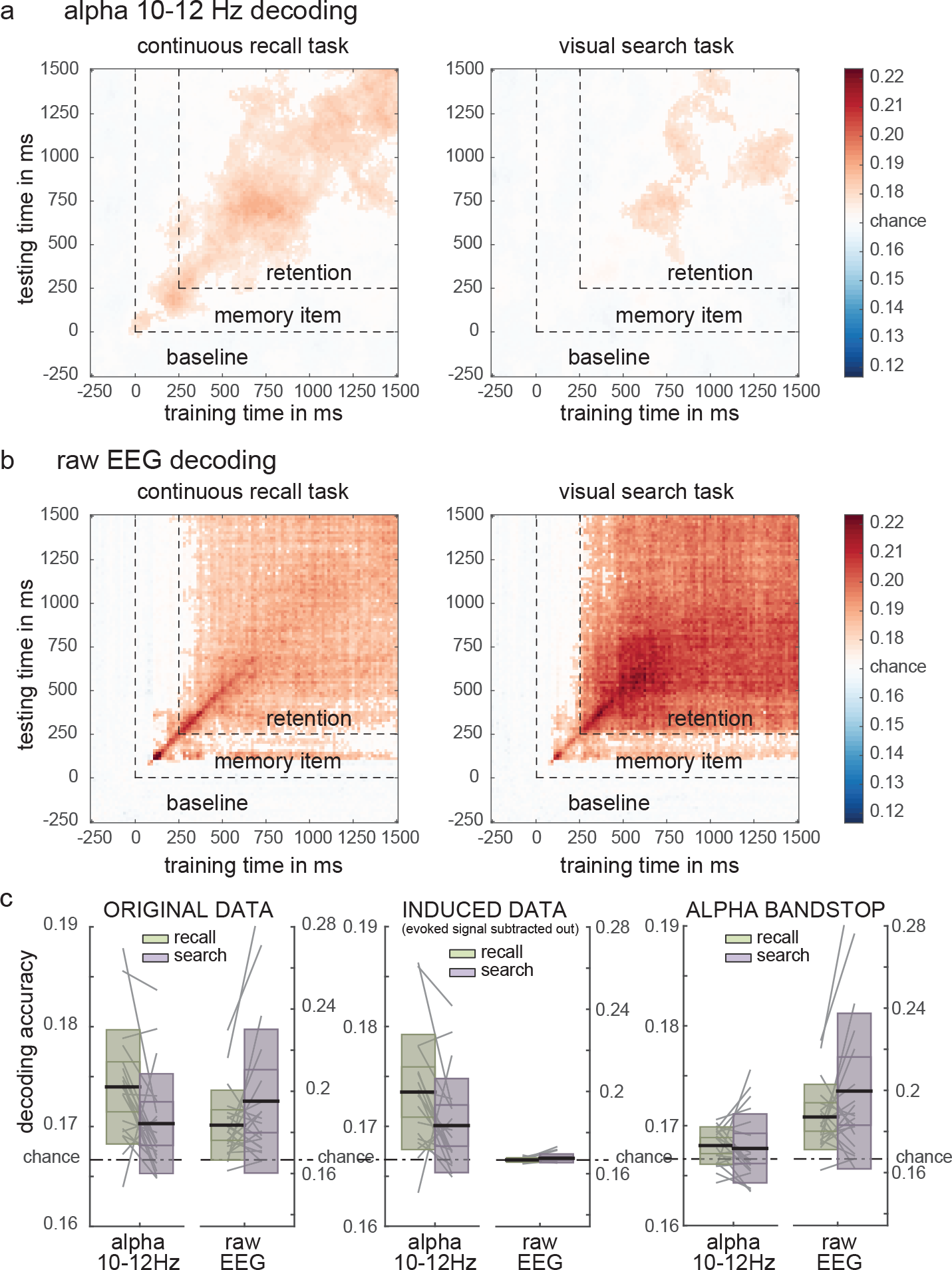
Goal-dependent WM traces contingent on neural measure. **(a)** Generalization Across Time (GAT) decoding of orientation using alpha-power (10-12 Hz) in both tasks. **(b)** GAT decoding of orientation using raw EEG in both tasks. Saturated colors survived cluster based permutation testing at *P*<.05. **(c)** The task-measure interaction using all data (left panel), using induced data, where the ERP is subtracted out (middle panel), and using alpha-bandstopped data (right panel). The thick horizontal black stripes indicate mean classification accuracy. The y-limits of the boxes around the black stripes respectively indicate the 95% confidence interval (closest to the mean) and the standard deviation (furthest away from the mean). Grey stripes in the background indicate single subject values.

Both raw EEG and alpha revealed solid orientation decoding in the retention interval in both tasks. Raw EEG decoding generalizes across the entire retention interval as evidenced by stable above chance decoding in nearly all train-test time combinations within the retention period. Alpha decoding was a little more dynamic with less off-diagonal decoding, but still shows clear above chance on-diagonal decoding in the continuous recall task. Importantly, the data show an interaction between measure and task type. An analysis of the average decoding accuracies in the retention interval of the GAT, using a 2 (task type: search/continuous) × 2 (measure: alpha/raw) repeated measures ANOVA (see Figure 2c, left panel) revealed a main effect of measure, showing that decoding is better using raw EEG than alpha-band (*F*_1,19_=11.61, *P*=0.003), no main effect of task, showing that memoranda can be decoded equally well in both tasks (*F*_1,19_=1.37, *P*=0.257) and an interaction between measure and task, showing that decoding was better in the alpha-band power range for the continuous recall task than for the search task, whereas decoding was better using raw EEG for the visual search task than for the continuous recall task (*F*_1,19_=7.13, *P*=0.015). Performing the analysis only on the diagonal of the GAT produced qualitatively and quantitatively equivalent results.

This interaction suggests that raw EEG signal and alpha-band power decoding have different root causes, and thus reflect dissociable maintenance mechanisms associated with different task goals. While both mechanisms are active during both tasks, raw EEG enabled more robust decoding in the search task, which encouraged the maintenance of the object itself, while alpha-band power enabled more robust decoding in the continuous recall task, which encouraged the maintenance of a location (i.e., the intersection point of the bar with the rim). Indeed, past work has established that the multivariate pattern of alpha-band power tracks locations held in spatial WM (20) and the locus of covert spatial attention (18, 19). It is unlikely that these effects are caused by leakage of the stimulus-encoding phase into the maintenance period. In the raw EEG signal, the relative strength of decoding accuracy in the search task superseded that of the continuous recall task only *after* the encoding period (see Supplementary Fig. S1b, right panel), while the alpha signal was unaffected by stimulus evoked activity altogether (see induced analyses below, including Supplementary Fig. S2).

However, to prove that the signals are truly dissociable requires one to demonstrate that they each carry unique information. Therefore, a second analysis approach showed that either signal could be removed, without the other being affected. We first abolished raw EEG decoding by subtracting the average evoked response from every trial, producing an induced (non-evoked) signal (see Supplementary Methods). This did not affect alpha-band decoding, while selectively eliminating above chance decoding of raw EEG (see Fig. 2c, middle panel and Supplementary Fig. S2). Conversely, when applying a bandstop filter to remove alpha-related activity, alpha-band decoding was virtually abolished, while raw EEG decoding survived (see Supplementary Methods, Fig. 2c, right panel and Supplementary Fig. S3). This confirms the ontological independence of the two signals.

A final analysis applied forward modeling to determine to what extent these two signals exhibited continuous graded channel tuning functions (CTFs, see Supplementary Methods), and whether the shapes of these CTFs were modulated by task goal. Fig. 3a and 3b show how the CTFs develop over time, for the alpha-band and raw EEG, separately for both tasks. Here too the pattern shows an interaction, reflecting a more sustained alpha-based CTF for the continuous recall task than the search task, while the reverse is true for the raw EEG CTF. A 2 (task type: search/continuous) × 2 (measure: alpha/raw) × 6 (stimulus orientation) ANOVA on average CTFs, averaging over all train-test combinations in the retention interval (see Fig. 3c and Supplementary Methods) confirmed the above results. There were main effects of measure (*F*_1,19_=103.07, *P*<10^-8^) and stimulus orientation (*F*_5,95_=84.26, *P*<10^-32^), but no main effect of task (*F*_1,19_=0.17, *P*=0.687) and no significant interactions between task and orientation (*F*_5,95_=0.10, *P*=0.993) or between measure and task (*F*_1,19_=0.16, *P*=0.694). However, there were strong interactions between measure and orientation (*F*_5,95_=8.65, *P*<10^-6^) and, most importantly, between task, measure and stimulus orientation (*F*_5,95_=3.99, *P*=0.003) confirming preferential orientation tuning in alpha during the continuous recall task and preferential orientation tuning in raw EEG during in the visual search task.

**Figure 3.**
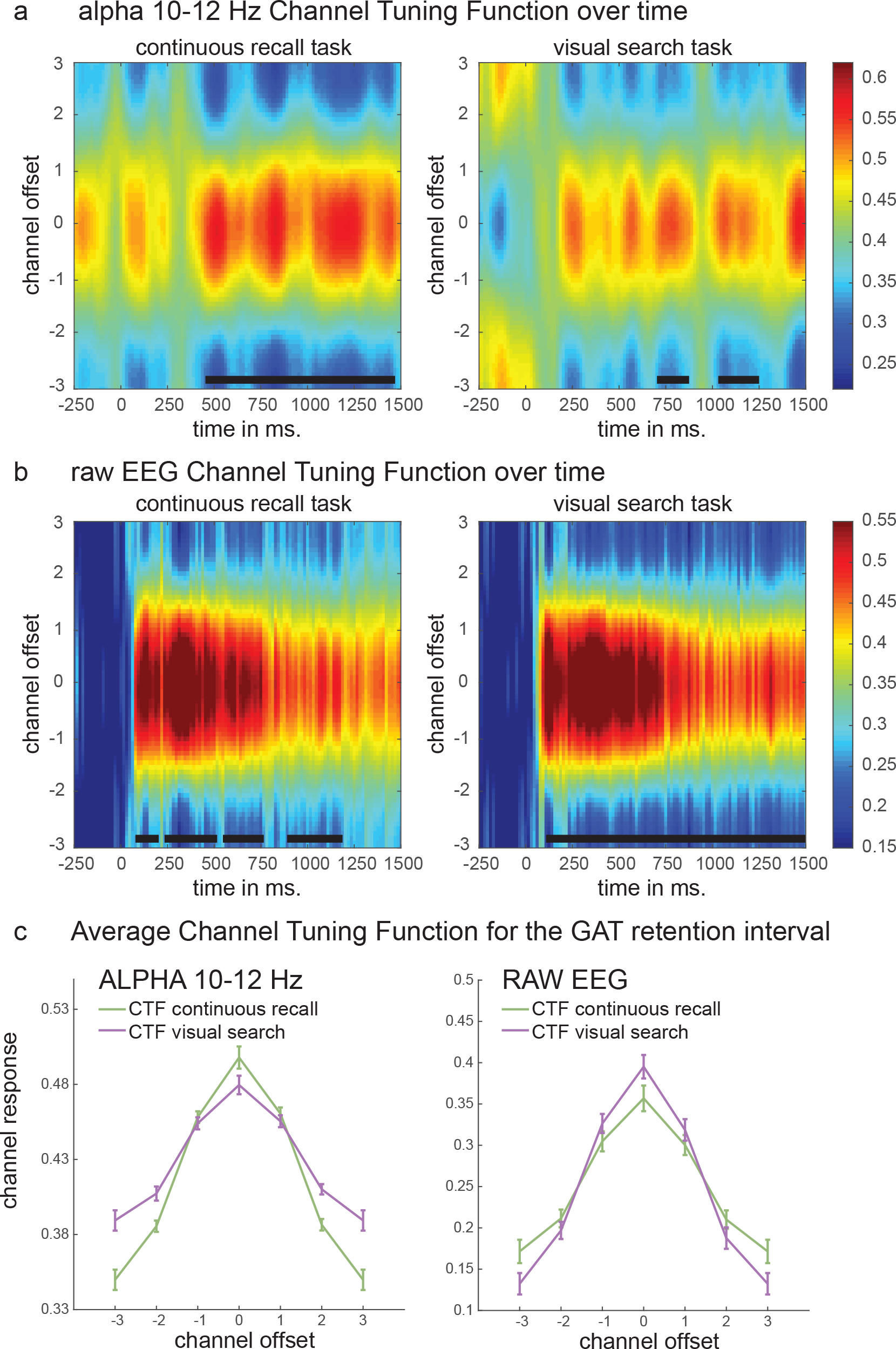
Channel tuning functions (CTF). **(a)** CTF over time based on alpha-band power (10-12 Hz). Black lines at the bottom indicated cluster-based significance of the slope of the CTF tested against 0. **(b)** CTF over time based on raw EEG. **(c)** Average CTFs over the entire train-test retention interval, for both tasks, separately for alpha-power (right) and raw EEG (left). These show a clear interaction.

Together, these data provide strong evidence for the existence of at least two dissociable goal-dependent maintenance mechanisms, either predominantly supporting a task favoring an object–based representation, or supporting one favoring a location-based representation. Orientation-dependent modulation of raw EEG signals seems to be dominant in tasks favoring an object-based representation, and seems to be expressed as a elevated firing signal in the EEG locked to the onset moment of retention (ERPs in Fig. 4), which is distinct from the oscillatory signal that is observed in the alpha range^also see (22)^ (Supplementary Fig. S3). Alpha-band on the other hand has a clear oscillatory signature that is not phase-locked to the onset moment of retention (Supplementary Fig. S2). We suggest that this reflects modulation of ongoing synchronous oscillatory activity related to spatial attention (18, 19) and possibly motor planning (23) associated with the spatial location of the planned response.

**Figure 4.**
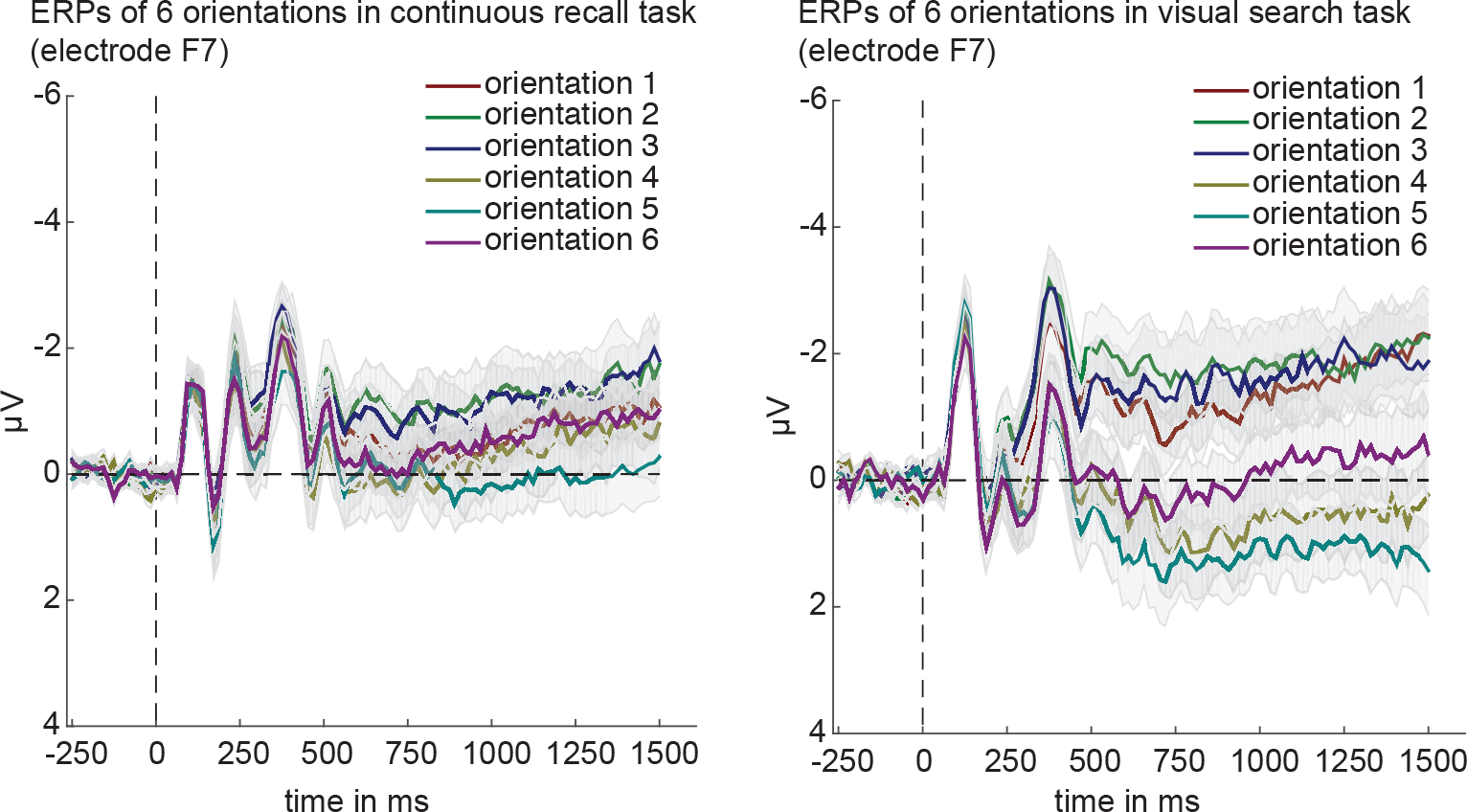
ERPs from the raw signal in electrode F7 for the 6 cardinal orientations in the two tasks. Left: continuous recall task. Right: visual search task. ERPs of the continuous recall task showed similar but weaker modulation by orientation compared to the visual search task, plausibly due to the fact that an alpha-related maintenance mechanism prevailed in the continuous recall task.

